# Modulation of Early Mitotic Inhibitor 1 (EMI1) Depletion on the Sensitivity of PARP Inhibitors in BRCA1 Mutated Triple-Negative Breast Cancer Cells

**DOI:** 10.1101/2020.06.09.142026

**Authors:** Dina Moustafa, Maha R. Abd Elwahed, Hanaa H. Elsaid, Jeffrey D. Parvin

## Abstract

Triple negative breast cancer (TNBC) represents approximately 10–15% of all breast cancers and has a poor outcome as it lacks a receptor target for therapy, and TNBC is frequently associated with a germline mutation of *BRCA1*. Poly (ADP-ribose) polymerase inhibitor (PARPi) drugs have demonstrated some effectiveness in treating *BRCA1* or *BRCA2* mutated breast and ovarian cancers but resistance to PARPi is common. Published results found that resistance to Olaparib, a PARPi, can be due to downregulation of EMI1 and the consequent upregulation of the RAD51 recombinase. Using a tissue culture-based cell viability assay, we extended those observations to another PARPi and to other chemotherapy drugs that affect DNA repair or the cell cycle. As we expected, EMI1 downregulation resulted in resistance to another PARPi drug, Talazoparib. EMI1 downregulation also led to resistance to other cytotoxic drugs, Cisplatin and CHK1 inhibitor. Surprisingly, EMI1 depletion also led to resistance to a MEK inhibitor, though this inhibitor blocks cells in G1 phase of the cell cycle and would not be expected to be sensitive to EMI1 levels. Notably, increasing the RAD51 protein expression only partially recapitulated the effects of EMI1 depletion in causing resistance to different PARPi and the other cytotoxic drugs. These results suggest that the downstream effects of EMI1 downregulation that contribute to PARPi resistance are increasing the concentration of RAD51 protein in the cell and blocking mitotic entry. We found that combining CHK1 inhibitor with olaparib results in restoration of sensitivity even when EMI1 expression is downregulated. This combination therapy may be a means to overcome the PARPi resistance in BRCA1-deficient TNBC cells.

## Introduction

The prognosis of breast cancer depends on several characteristic features, namely, estrogen receptor (ER), progesterone receptor (PR), and HER2 receptor expression and mutation status. The phenotype of germline *BRCA1* mutations are usually characterized by aggressiveness, high grade, and are more likely to be triple-negative (ER-, PR-, and HER2-) (1,2). *BRCA1* or *BRCA2* mutant breast tumor cells are deficient in the repair of DNA double strand breaks (DSB) via the homology directed repair (HDR) mechanism. This repair mechanism is sequence conserving, and cells lacking the appropriate function of either of these genes have an increased rate of mutation. PARPi block the base excision repair mechanism for single strand breaks, but PARPi in combination with a *BRCA1* or *BRCA2* mutation are synthetic lethal, and such cells are sensitive to PARPi (3,4). Thus, tumors deficient in BRCA1 or BRCA2 are highly vulnerable to the effects of PARP inhibition (2,5). PARP inhibitors have shown promise in cancer therapy via a mechanism dependent on synthetic lethality; the inhibition of PARP results in the accumulation of a significant amount of double-strand breaks (DSB) by interfering with replication fork progression at the site of DNA damage (6). Normal cells with a *BRCA* mutation are often heterozygous for the abnormal gene, are able to repair the DSB and survive unaffected under conditions of PARP inhibition. Contrariwise, cancer cells may retain only the mutated copy of *BRCA1* or *BRCA2*, conferring inadequate DNA repair and susceptibility to the synthetic lethality of PARPi (2,7). Despite the promising preliminary results, prolonged treatment of breast or ovarian cancer with PARPi is frequently associated with acquired resistance to this therapy. There are multiple mechanisms of resistance to PARPi chemotherapy, some of them are targetable for therapy including the modulation of EMI1 and RAD51 expression levels (8,9).

HDR is a complex pathway that requires not only the efficient use of BRCA1 and BRCA2 proteins but a number of other related proteins including, among others, REV7, PTIP, RIF1 and RAD51 (10) The key protein in HDR is the recombinase, RAD51; the activity and level of the RAD51 protein is regulated in part by p53 (11,12). The level of RAD51 is biologically important in regulating DNA repair(13) and RAD51 overexpression, is associated with an increase in the spontaneous level of HDR with subsequent resistance to ionizing radiation (14).

Another mechanism for resistance to DNA damage in actively dividing cells is the activation of distinct cell checkpoint responses, which pause the cell from progressing to the next cell cycle, enabling DNA repair and promoting cell survival. Upon DNA damage, G2 arrest is triggered by p21-dependent EMI1 downregulation(15). In human cells, the EMI1 protein level is uniform throughout the cell cycle except for its absence from mitotic entry to G1. EMI1 restrains the activation of APC in interphase and its degradation in prophase by F-box protein β-Trcp1 leads to the activation of APC (16,17). Thus, EMI1 down-regulation activates APC, which in turn degrades key substrates that prevent the G2-arrested cells from entering mitosis (15). Additively PARP1 deficient cells show a stronger G2 checkpoint response, which is regulated by the ATR/ CHK1 pathway(18,19). Combination of PARPi with ATRi leads to complete ovarian tumor regression in an HDR-deficient PDX model(19)

Published experiments (20) demonstrated that downregulation of EMI1 in BRCA1-deficient breast cancer cells led to a decrease in sensitivity of the cells to PARPi. They suggested that, in the absence of DNA damage, the E3 ubiquitin ligase activity of EMI1 regulates the stability of RAD51. Marzio et al showed that the resistance to PARPi conferred by decreased EMI1 concentration could be replicated by over-expressing RAD51. Marzio et al tested only olaparib as the PARPi and they suggested, but did not test, that the resistance conferred by low EMI1 concentration could be reversed by including a CHK1 inhibitor to their cell system. In the current study, we tested a second PARPi as well as other DNA-damaging or cytostatic drugs for resistance in cells with low EMI1 concentration, and we tested whether combining CHK1 inhibitor with the PARPi could rescue sensitivity in these cells even though they had low EMI1 concentrations.

## Materials and Methods

### Cell culture

Cell line MDA MB 436 was propagated in DMEM media supplemented with 10% FBS (VWR Seradigm Life Science), 1% L-glutamine and 1% sodium pyruvate (Life Technology-Thermo-Fisher). MDA MB 231 was propagated in DMEM media with 10% FBS, 1% L-glutamine and 1% sodium pyruvate and 1% penicillin/streptomycin. MCF7 propagated in DMEM media with 10% FBS, 1% L-glutamine and 1% sodium pyruvate, 1% penicillin/streptomycin and 0.01mg/ml bovine insulin. SUM149PT was propagated in Ham’s F12/L-glutamine media supplemented with 5% FBS,10mM HEPES, 5 µg/ml insulin and 1 µg/ml hydrocortisone. Cell lines were confirmed by STR fingerprinting and tested periodically for mycoplasma contamination using PCR kit and MycoAlert TM Mycoplasma Detection Kit (Catalog #: LT07-418) and the results were negative.

### Gene depletion by siRNA transfection

DNA and RNA oligonucleotides were purchased from IDT. Cells were transfected using either non-targeting siRNA (GL2) (38) that targets the luciferase gene and served as a negative control or siRNAs directed toward EMI1. (Sequences of siRNAs used in this study are provided in supplementary table 1).

### Plasmids

RAD51 cDNA (gift of R. Fishel, Ohio State University) was inserted into a pcDNA3 vector and empty pcDNA3 vector was used as a negative control. Transfections were done using either oligofectamine for siRNA transfection (Invitrogen P/N 58303) using 50 pmol of each siRNA or Lipofectamine 2000, Invitrogen P/N 52887) for RAD51 vector or Empty pcDNA3 vector, using 3 µg of each vector.

For MDA MB 436, 5×10^5^ cells were seeded per well in a 6 well plate, 24 h later transfections done, and after overnight incubation media was changed. 3 h later, a second transfection was applied. For MDA MB 231, 3×10^5^ cells were seeded per well of a 6 well plate, and a similar transfection protocol was applied.

### Immunoblotting

Protein was isolated using protein lysis buffer (50 mM Tris PH 7.9, 300 mM NaCl, 2.5% NP40, 1 mM EDTA, 5% glycerol). Then samples were resolved on 8% SDS-PAGE gels and transferred to PVDF membrane. The following primary antibodies were used according to the manufacturer’s recommendations: EMI1 3D2D6 (Thermo-Fisher (1:500)) or SAB2100793, (Sigma-Aldrich (1mg/ml)), BRCA1(1:500) (39), RAD51 (GeneTex, Cat. No. GTX70230 (1:500)). RNA Helicase A (RHA; 1:20,000) (39) (Sigma-Aldrich) was used as loading control. Secondary antibodies were used according to the manufacturer’s specification. anti-rabbit horseradish peroxidase (HRP) 1:5,000 (Cell Signaling Technology, cat. No. 7074), anti-mouse HRP 1:5,000 (Cell Signaling Technology, catalog no. 7076).

### Cell viability assay

Different cell lines were transfected as described above, and 24 h after the second transfection, cells were plated by 2500 cells/well for each condition in 96 well plates. 24h later drug is added according to each experiment design. After incubation 72 h for cisplatin incubation and 96 h for all other drugs, viability was measured using alamarBlue (Cat. No DAL1025, Thermo-Fisher Scientific) according to manufacturer specifications.

### Statistical Analysis

Data are representative of at least three biologically independent experiments. All datasets were analyzed by Student’s paired t test. Results are indicated as statistically significant if p<0.05 (*) or p<0.01 (**).

## Results

### Reduction EMI1 expression and resistance to PARPi, cisplatin, and CHKi

As a first step in extending the findings of Marzio et al, we compared two cell lines, *BRCA1*-mutated MDA-MB-436 to *BRCA1*-wild-type MDA-MB-231 cells. Both cell lines recapitulate the phenotypes of Triple Negative Breast Cancer (TNBC) cells. We found, consistent with expectations, that the MDA-MB-436 cells were much more sensitive to olaparib; at all datapoints from 2.5 µM to 80 µM olaparib there was a statistically significant difference in sensitivity to the drug. The IC50 was about 2.5 µM olaparib in MDA-MB-436 cells and about 80 µM olaparib in the *BRCA1*-wild-type MDA-MB-231 (Figure 1A, *left*). Similarly, the MDA-MB-436 cells were more sensitive to cisplatin than were MDA-MB-231 cells, though the change in IC50 was not as dramatic (Figure 1A, *right*). Consistent with the Marzio et al report, depletion of EMI1 in *BRCA1* mutated MDA-MB-436 cells by transfection with either of two siRNAs reversed the sensitivity to olaparib and shifted the IC50 in this experiment from about 1.25 µM with normal EMI1 to about 40 µM when EMI1 was depleted (Figure 1B). Western blots revealed that the depletion of EMI1 with each siRNA was effective and resulted in an increase in the abundance of RAD51 protein. Immunoblots of RNA Helicase A (RHA) were used as a loading control.

**Figure 1.**
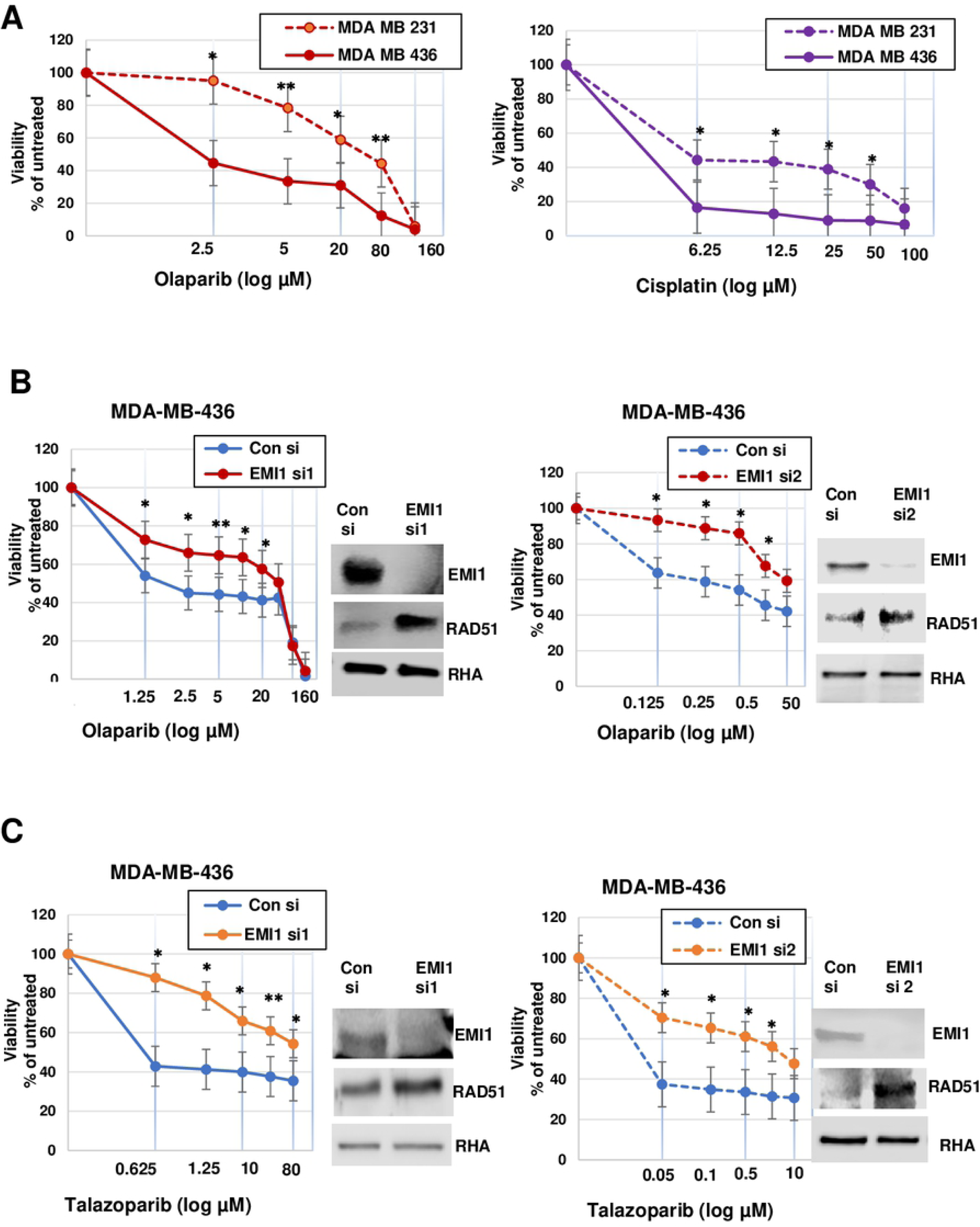
Depletion of EMI1 in BRCA1 mutant MDA-MB-436 cells confers resistance to PARPi. **(A)** Cell lines that mimic TNBC, MDA MB 436 cells (*BRCA1* mutant) and MDA MB 231 cells (expressing wild-type BRCA1) were grown in medium containing olaparib (*left*) or cisplatin (*right*) at the indicated concentrations for 96 h for olaparib or 72 h for cisplatin when cells were assayed for proliferation. For each transfection, the proliferation in the absence of drug was set at 100%. The x-axis is in log_10_ and the datapoints from samples containing no drug (vehicle only) was placed on the y-intercept. Results represent the mean and SEM of three replicate experiments. Datapoints at each concentration of olaparib were analyzed by the two-sided student’s t test, and * indicates p<0.05, and ** indicates p<0.01. **(B)** MDA MB 436 cells were transfected with a control siRNA or EMI1-specific siRNA1 (*left*) or control siRNA or EMI1 siRNA2 (*right*) and then grown in medium containing olaparib at the indicated concentrations for 96 h when cells were assayed for proliferation. Western blots are shown from one of the replicates used in the proliferation assay and stained for protein abundance of EMI1, RAD51, and RNA helicase A (RHA), as indicated. Results represent the mean and SEM of five replicate experiments testing EMI1 specific siRNA1 and four replicates for experiments testing siRNA2. **(C)** MDA-MB-436 cells were transfected as in panel B, and subjected to the indicated concentrations of talazoparib in the culture medium for 96 h when proliferation and protein expression was measured as described above. Results represent the mean and SEM of three replicate experiments.

Extending the results from Marzio et al to a second PARPi, we found that depletion of EMI1 by transfection of either of the two siRNAs from MDA-MB-436 cells rendered the cells resistant to talazoparib and the IC50 changed from less than 0.05 µM with normal EMI1 levels to about 10 µM with EMI1 levels reduced by transfection of specific siRNA (Figure 1C), consistent with the expectation of PARPi resistance when the EMI1 levels were depleted.

Strikingly, we found that low EMI1 levels in these *BRCA1* mutated MDA-MB-436 cells can modulate the sensitivity to other cytotoxic drugs, such as cisplatin. Results in Figure 2A show a significant upward shift of the IC50 of cisplatin from lower than 0.5 µM with normal EMI1 levels to about 5 µM with EMI1 levels reduced by specific siRNA transfection. Since the titration of cisplatin when testing the sensitivity of MDA-MB-436 cells transfected with EMI1-specific siRNA-1 were near the lowest levels at the lowest dose tested, 5 µM (Figure 2A, *left*), we tested lower concentrations of cisplatin for EMI1-specific siRNA-2 (Figure 2A, *right*). We found that the lowest dose, 0.5 µM was higher than the IC50, but in control siRNA transfected cells the IC50 was between 1.5 µM and 10 µM cisplatin.

**Figure 2.**
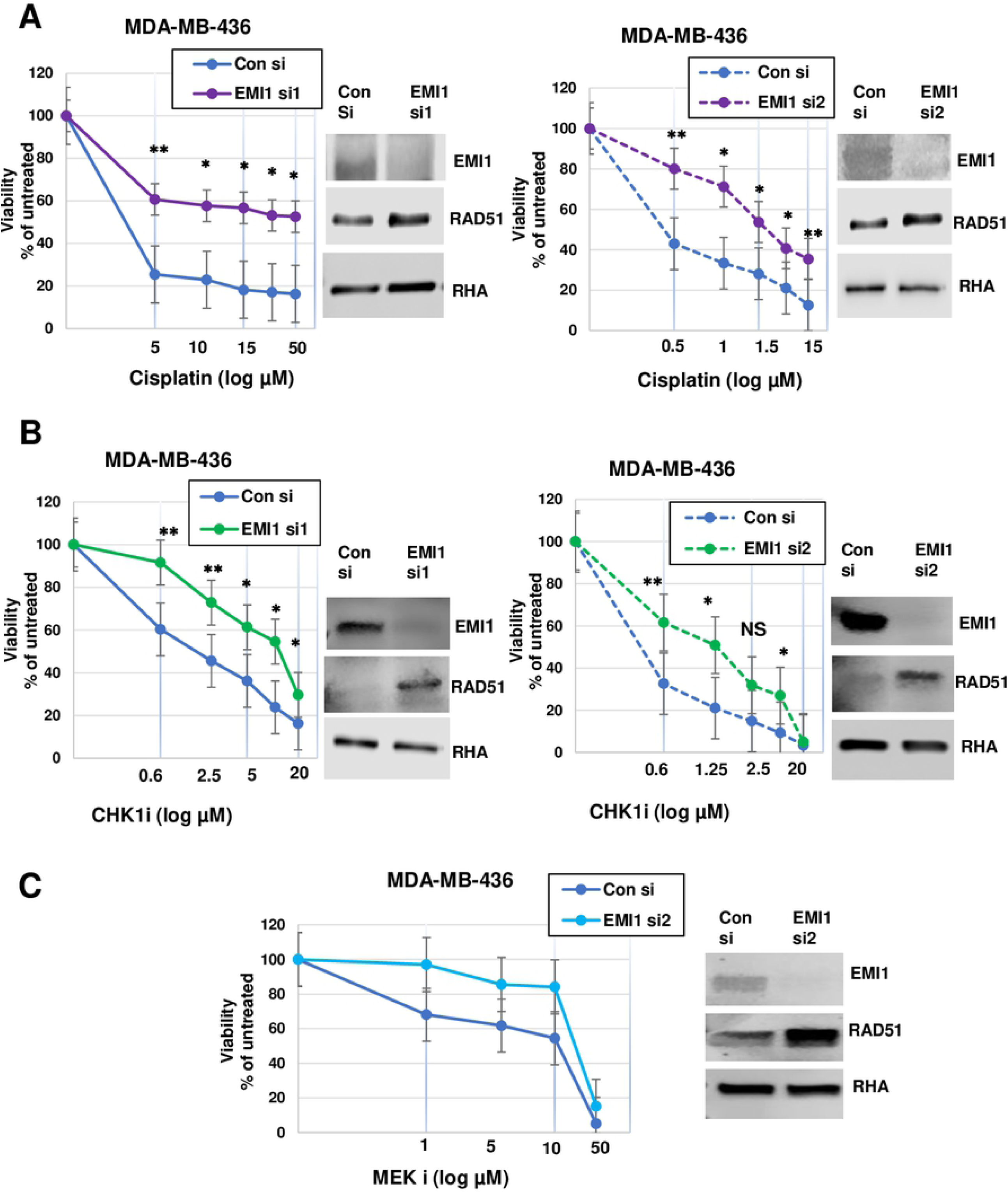
Depletion of EMI1 in BRCA1 mutant MDA-MB-436 cells confers resistance to cytostatic drugs. **(A)** MDA-MB-436 cells, transfected as in Figure 1, were subjected to the indicated concentrations of cisplatin for 72 h followed by a proliferation assay and an immunoblot of one of the replicates. The x-axis is in log_10_ and the datapoints from samples containing no drug (vehicle only) were placed on the y-intercept. Results represent the mean and SEM of three replicate experiments. Datapoints at each concentration of cisplatin were analyzed by the two-sided students t test, and * indicates p<0.05, and ** indicates p<0.01. **(B)** MDA-MB-436 cells were transfected as in Figure 1 and subjected to the CHK1 inhibitor (SB218078) in the growth medium for 96 h and analyzed as described above. Results represent the mean and SEM of three replicate experiments. **(C)** MDA-MB-436 cells were transfected with control siRNA and EMI1-specific siRNA2, followed by inclusion of the MEKi (selumetinib) in culture medium at the indicated concentrations for 96 h, followed by analysis for proliferation and protein abundance as described above. Results represent the mean and SEM of three replicate experiments.

We also found that reduction of EMI1 levels resulted in a decrease in sensitivity of CHK1 inhibitor (SB218078) when used as a monotherapy. The IC50 for the CHKi was about 2.5 µM for the control siRNA transfected MDA-MB-436 cells, and transfection of EMI1-specific siRNA-1 resulted in a relative resistance to the CHKi, with an IC50 of about 15 µM CHKi (Figure 2B *left*). Repeat of this experiment using EMI1-specific siRNA-2 revealed a similar trend, though there was a minor change in the IC50 for both control and EMI1-specific siRNAs.

Interestingly, when we tested the effect of depleted EMI1 levels toward another cytostatic drug, MEK inhibitor (selumetinib AZD6244), we found a small shift in sensitivity by a reduction in the EMI1 protein levels. The minor change in sensitivity was not statistically significant, and perhaps reflects that depletion of EMI1 is known to arrest cells in mitosis (15) and MEKi blocks cells in G1 phase of the cell cycle (21,22).

We tested a second *BRCA1* mutated cell line, SUM149PT. We found that depletion of EMI1 by siRNA transfection diminished the sensitivity of each, olaparib and CHK1 inhibitor. RNAi depletion of EMI1, using ether of the two specific siRNAs in this cell line, was effective in reducing the expression of EMI1 and with the downstream effect of increasing RAD51 protein levels (Figure 3C). The effect of EMI1 depletion on IC50 was shifted from 5 µM to about more than 10 µM olaparib (Figure 3A) and from 0.6 µM to 1.25 µM CHK1i (Figure 3B). Although the effect of EMI1 depletion on resistance to these two drugs was not as dramatic as observed with MDA-MB-436 cells, the results were statistically significant at most concentrations of olaparib and at some concentrations of CHK1i, as indicated in the Figure. The trends in these observations using SUM-149PT cells were consistent with our previous findings using MDA-MB-436 cells and with published results (20). Taken together with our other results, these results suggest that low expression of EMI1 can modulate the sensitivity to several chemotherapeutic drugs, not only PARPi, and this is a potential mechanism to cancer therapy resistance.

**Figure 3.**
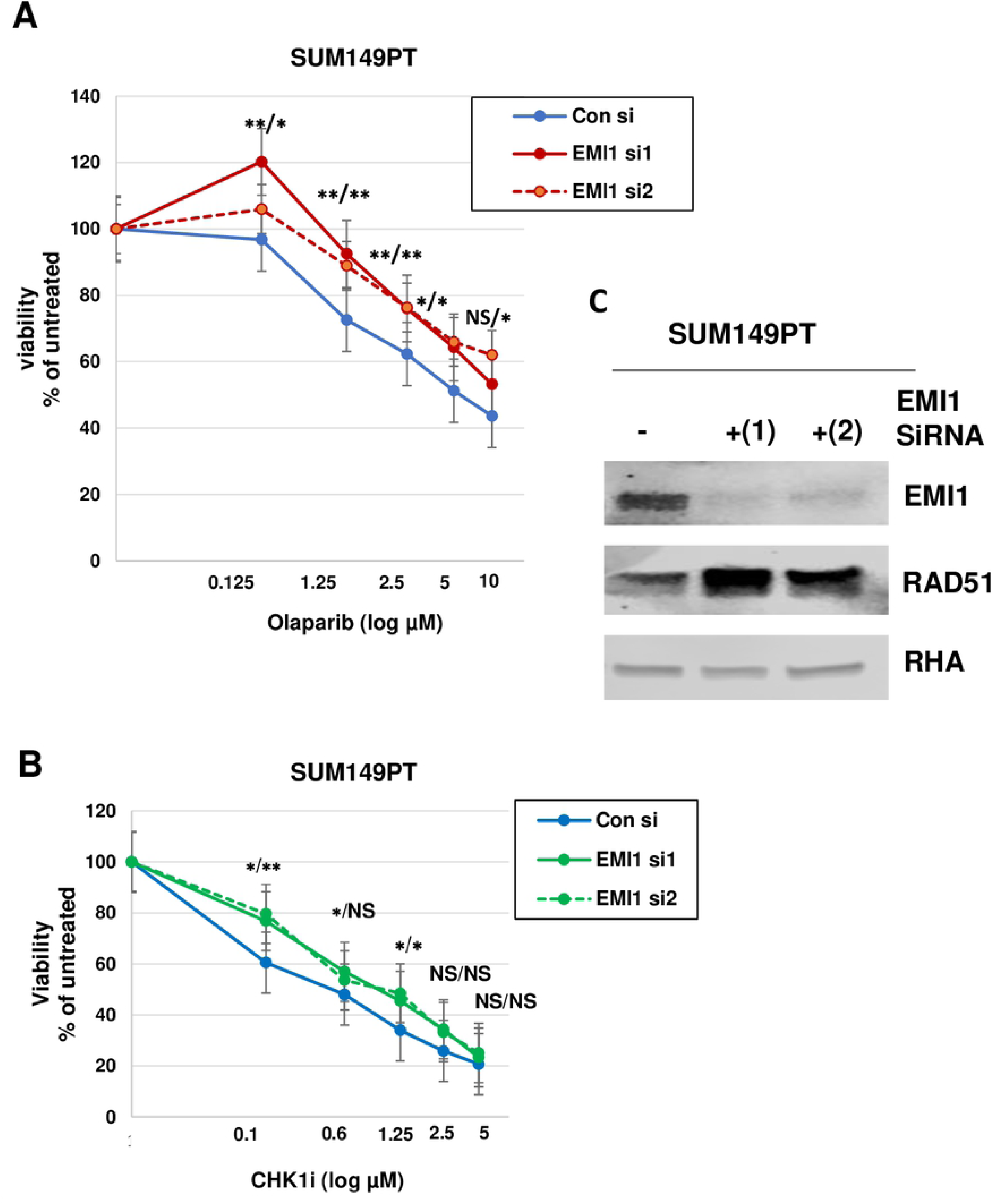
Depletion of EMI1 in BRCA1 mutant SUM149PT cells confers resistance to olaparib and CHK1i. **(A)** SUM149PT cells were transfected with control siRNA, EMI1-specific siRNA1 or EMI1-specific siRNA2, followed by inclusion of olaparib in culture medium at the indicated concentrations for 96 h. Cells were then analyzed for proliferation, and the measure in samples without drug (vehicle only) were set at 100%. Results represent the mean and SEM of three replicate experiments. Datapoints at each concentration of olaparib were analyzed by the two-sided students t test, and * represents p<0.05, ** represents p<0.01, and NS represents p>0.05. At each concentration of drug, the first asterisks represent EMI siRNA1 versus control, and the second asterisks represent EMI1 siRNA2 versus control. Results represent the mean and SEM of three replicate experiments. **(B)** SUM149PT cells were transfected as in panel A and subjected to growth in medium with CHK1i (SB218078) at the indicated concentrations for 96 h followed by a proliferation assay. Results were analyzed as in panel A and represent the mean and SEM of three replicate experiments. **(C)** Protein lysates from one of the replicates used in panels A and B were analyzed by immunoblot and probed for EMI1, RAD51, and RHA protein abundance, as indicated.

### Effect of EMI1 depletion on resistance to some chemotherapy drugs is not phenocopied by elevated RAD51 protein abundance

It had been observed that resistance to olaparib due to EMI1 depletion could be recapitulated by expressing RAD51 at higher levels (20). The results of that study indicated that RAD51 protein was a substrate of the EMI1 ubiquitin ligase, and this regulatory event was the key downstream effect of low EMI1 levels. We tested the effect of RAD51 overexpression in BRCA1-mutant cells on sensitivity to different cytotoxic drugs including the PARPi. We prepared an expression vector for human RAD51 that can be used for transfection of mammalian cells and increase the expression of RAD51 protein in BRCA1-mutant MDA-MB-436 cells. As measured by immunoblot analysis, transient transfection of RAD51 in MDA-MB-436 cells resulted in a several-fold increase in RAD51 protein levels (Figure 4A, *right*). Similar to the results of Marzio et al, we found that the resistance to olaparib conferred by EMI1 depletion was largely recapitulated by RAD51 overexpression (Figure 4A, *left*). The IC50 for olaparib was shifted from lower than 0.125 µM with normal, endogenous RAD51 levels to about 0.25 µM with overexpressed RAD51 and, though the change was less dramatic than observed with EMI1 depletion (Figure 1B), it was statistically significant at each concentration of olaparib (Figure 4A *left*). By contrast, overexpression of RAD51 did shift the curve of the cells resistant to talazoparib, but for none of the data points were the differences statistically significant (Figure 4A, *right*). Similarly, overexpression of RAD51 resulted in a minor, and not statistically significant, shift of the curve for MDA-MBD-436 cells treated with cisplatin (Figure 4B *left*), CHK1 inhibitor (Figure 4B *right*) and MEK inhibitor (Figure 4C).These results indicated that high abundance of RAD51 protein, alone, cannot explain the resistance to these drugs by low expression of EMI1. Inactivation of EMI1 leads to premature APC/C activation in G2-phase, and consequently blocks cell entry into mitosis (23,24), and as expected, there was a G2 block evident in EMI1-depleted MDA-MB-436 cells (data not shown). PARP-induced cytotoxicity has been attributed to repeated cycles of both replication and mitosis showing that forced mitotic bypass through EMI1 depletion could largely rescue viability of HDR-deficient cells upon PARP inhibition (25). Taken together, we infer that two effects of EMI1 depletion, the increased abundance of RAD51 protein and a cell cycle block in G2, combine to overcome the effects of PARPi, cisplatin, and CHK1i.

**Figure 4.**
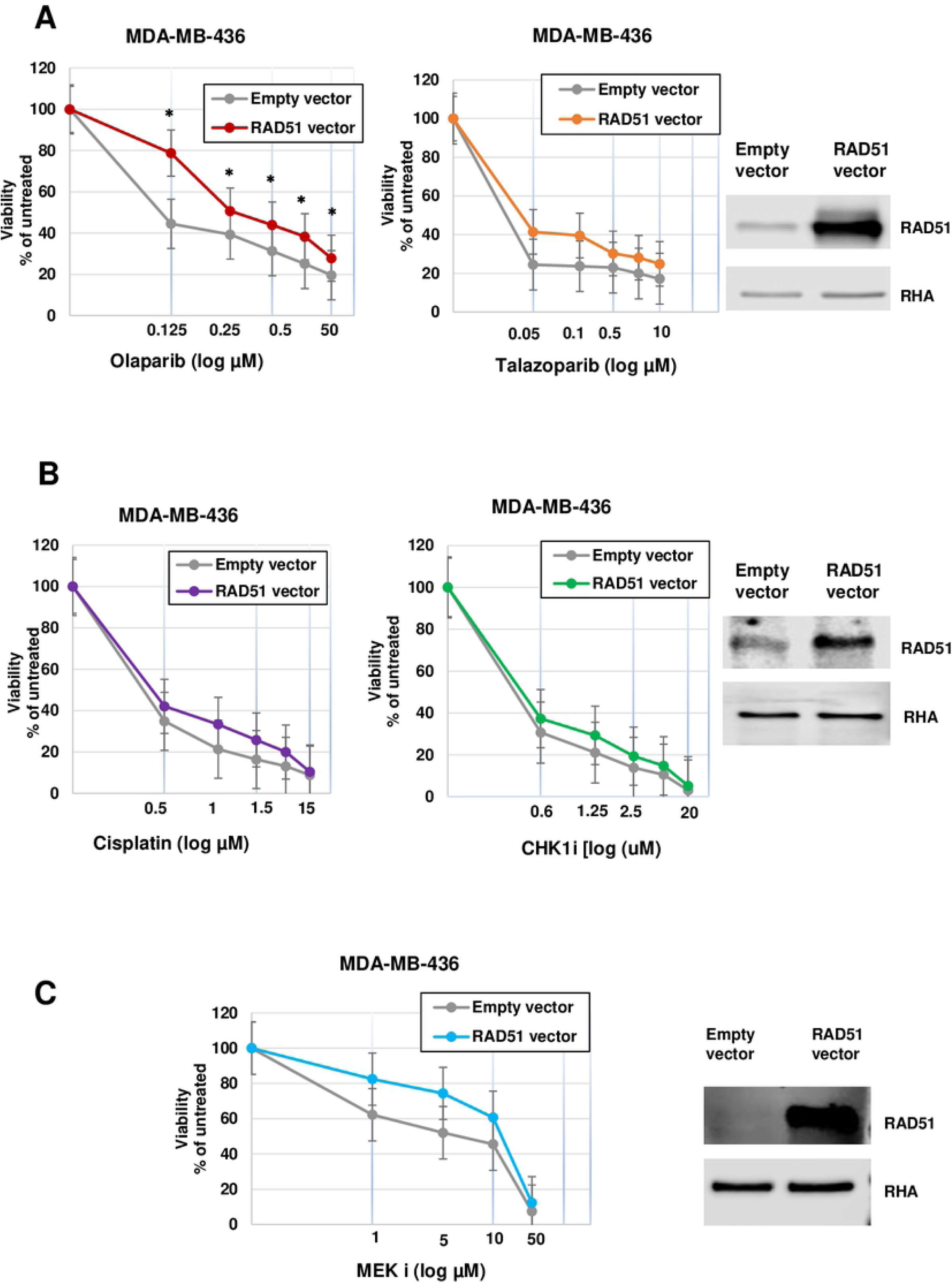
RAD51 overexpression only partially recapitulates effects of EMI1 depletion on resistance to cytostatic drugs in MDA-MB-436 cells. **(A)** MDA-MB-436 cells were transfected with pcDNA3 empty vector or with the same vector with the human RAD51 gene. Cells were grown for 96 h in medium containing the indicated concentration of olaparib (*left*) or talazoparib (*middle*) followed by a proliferation assay. Results represent the mean and SEM of three replicate experiments. Datapoints at each concentration of drug were analyzed by the two-sided student’s t test, and * represents p<0.05. Protein lysates from one replicate were analyzed by immunoblot for RAD51 protein abundance (*right*). **(B)** MDA-MB-436 cells were transfected as in panel A, and cells were grown in medium containing the indicated concentration of cisplatin (*left*) for 72 h or CHK1i (*middle*) for 96 h followed by a proliferation assay. Results were analyzed as in panel A and represent the mean and SEM of three replicate experiments. Protein lysates from one of the replicates were analyzed by immunoblot for RAD51 protein abundance (*right*). **(C)** MDA-MB-436 cells were transfected as in panel A, and cells were grown in medium containing the indicated concentration of MEKi (*left*) for 96 h followed by a proliferation assay. Results were analyzed as in panel A and represent the mean and SEM of three replicate experiments. Protein lysates from one of the replicates were analyzed by immunoblot for RAD51 protein abundance (*right*).

### Combining CHK1 inhibitor with PARPi restores sensitivity to BRCA1 mutated breast cancer cells with low EMI1 expression

In their earlier publication, Marzio and his colleagues suggested, but did not test, that the resistance to olaparib conferred by low EMI1 concentration could be reversed by including a CHK1 inhibitor (20). In addition, it was reported that a stronger DNA damage-induced G2 checkpoint dependent on CHK1 activation in the absence of PARP1 could be abolished by CHK1 siRNA, sensitizing PARP1-/- cells to IR-induced killing (18). Taking into consideration these results, we tested the combination of CHK1 inhibitor and olaparib in *BRCA1*-mutated MDA-MB-436 and SUM149PT cells. MDA-MB-436 samples were subjected to 10 µM CHK1i, and the olaparib was titrated from 0 to 160 µM, while, SUM149PT samples were subjected to 0.1 µM CHK1i, and the olaparib was titrated from 0 to 10 µM. In the presence of both, CHK1i and olaparib, the resistance due to EMI1 depletion was reversed (Figure 5A). Consistent with our results from Figure 2, in the presence of one drug, olaparib, EMI1 depletion resulted in a significantly higher percentage of viable cells than control depletion (solid lines in Figure 5A). By contrast, inclusion in culture media of CHK1i resulted in a sensitivity of these cells to olaparib in both control and EMI1-depleted cells. The IC50 for olaparib in EMI1 depleted cells changed from 40 µM to less than 2.5 µM when CHK1i was included in the culture medium, and the differences in proliferation under these conditions with two drugs was not statistically significant (p> 0.05). Similarly, when testing the combination of olaparib and CHK1i in SUM149PT cells, we found that the combination rendered EMI1-depleted cells sensitive (Figure 5B). Just as in the MDA-MB-436 cells, SUM149PT cells depleted of EMI1 were relatively resistant to olaparib, and the proliferation of these cells dropped in the presence of both drugs. When cells were transfected with EMI1-specific siRNA1, there was essentially no difference from the control siRNA transfected cells. Results were similar when depleting EMI1 from these cells using siRNA2; the IC50 for olaparib in EMI1-siRNA2 depleted cells was greater than 10 µM and in the presence of both drugs was about 2 µM. Though the magnitude of the change in sensitivity to two drugs when transfecting with siRNA2 was not as large as with siRNA1, the results were consistent. The p-values for student’s t-test comparisons are given in the legend to Figure 5B. These results suggest that CHK1 inhibitor in combination with PARPi could be a viable treatment option in these tumor types when EMI1 depletion causes resistance.

**Figure 5.**
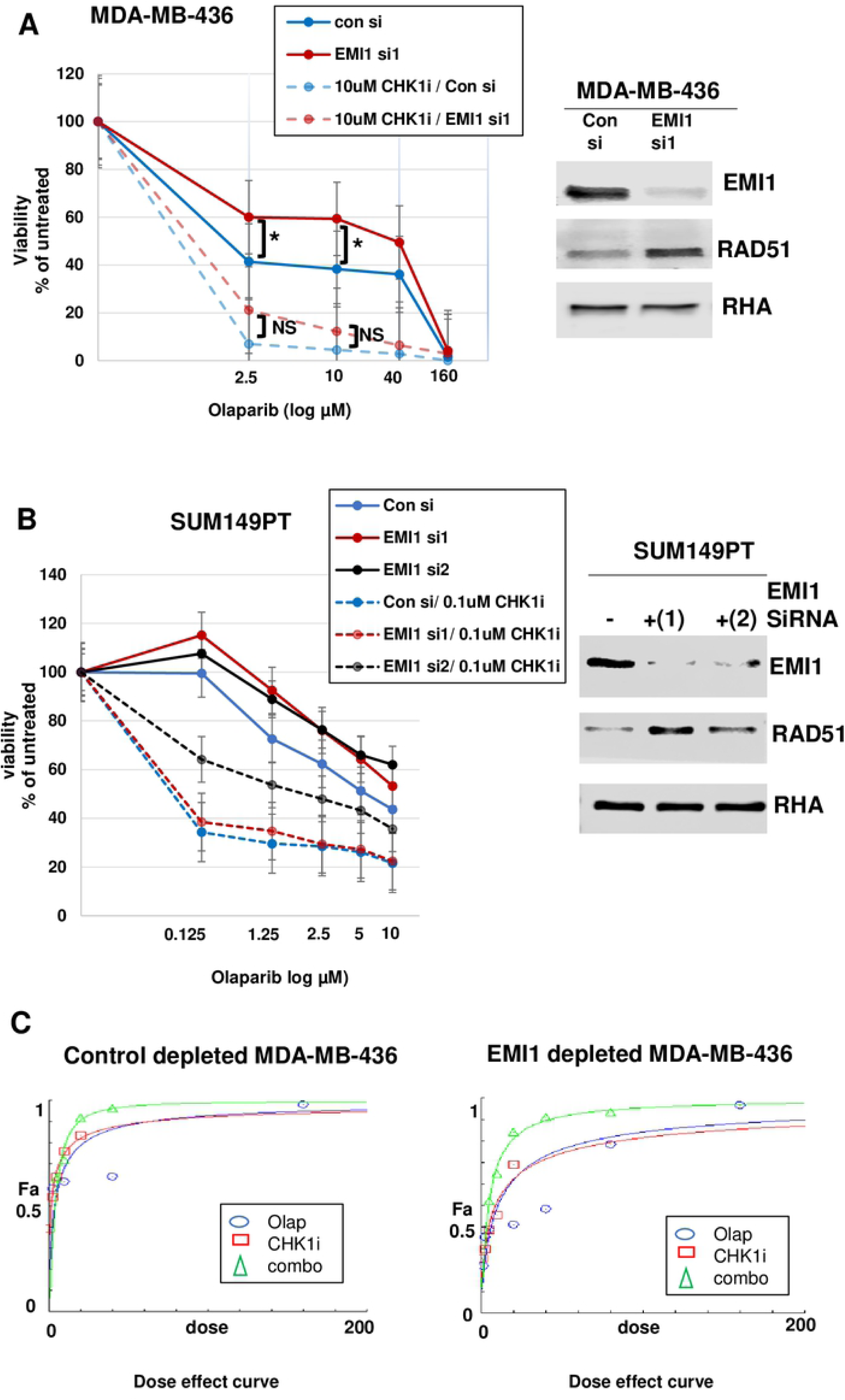
Combination of olaparib and CHK1i restores sensitivity to EMI1-depleted MDA-MB-436 and SUM149PT cells. **(A)** MDA-MB-436 cells were transfected with control siRNA or EMI1-specific siRNA1 and grown for 96 h in medium containing vehicle plus olaparib at the indicated concentrations (solid lines) or in medium containing 10 µM CHK1i plus olaparib at the indicated concentrations (dashed lines). Proliferation assays (n=3) measured the growth of the cells under each condition. The growth of control siRNA transfected cells was compared to the growth of EMI1 depleted cultures grown in the absence of CHK1i and in the presence of CHK1i. P-values comparing the growth of EMI1-depleted cells versus control siRNA-transfected cells were less than 0.05 at olaparib concentrations of 2.5 µM and 10 µM in the presence of a single drug, but in the presence of both drugs, the p-values were not significant. Protein lysates from one replicate were analyzed by immunoblot for EMI1 and RAD51 protein abundance (*right*). **(B)** SUM149PT cells were transfected with control siRNA or EMI1-specific siRNA1 or siRNA2 and grown for 96 h in medium containing vehicle plus olaparib at the indicated concentrations (solid lines) or in medium containing 0.1µM CHK1i plus olaparaib at the indicated concentrations (dashed lines) and analyzed as in panel A. When comparing EMI1-siRNA1 to control siRNA, p-values were less than 0.05 at concentrations of olaparib of 0.125 µM, 1.25 µM, 2.5 µM, and 5 µM when testing the single drug, and was not significant (p > 0.05) at all concentrations when comparing EMI1 siRNA1 to control siRNA in two drugs. When comparing EMI1-siRNA2 to control siRNA, p-values were less than 0.05 at all concentrations of olaparib tested in the single drug and in the two drug conditions. (C) CompuSyn software analysis of additive effects of olaparib and CHK1i in MDA-MB-436 cells. The dose effect (x-axis) was plotted against the fraction of the total drug effect (Fa, y-axis) for olaparib alone (blue ovals), CHK1i alone (red rectangles) or the combination of both drugs (green triangles) in control siRNA transfected cells (left) and EMI1 siRNA1 depleted cells (right).

We analyzed if the CHK1i and olaparib were acting synergistically in MDA-MD-436 cells by applying the Chou-Talalay method (26) and using the computer software CompuSyn. We observed a synergistic effect of the two drugs increased in the EMI1-depleted MDA-MB-436 cells (Figure 5C). This effect is evident in the increase in separation of the curve for the drug combination from the curves for the single drugs. Repeating the analysis using SUM149PT cell data also indicated synergism in the EMI1-depleted cells (data not shown).

## Discussion

In this study, we discovered: 1) depletion of EMI1 made *BRCA1*-mutant cells resistant to talazaparib, cisplatin, and CHK1i in addition to the previously shown resistance to olaparib (20); 2) the effect of depleting EMI1 on drug resistance was only partially recapitulated by over expression of RAD51, suggesting that EMI1 effects on cell cycle progression also contribute to the PARPi resistance; 3) combinations of EMI1 depletion and CHK1i restored sensitivity of the BRCA1-mutant cells toward PARPi; and 4) olaparib and CHK1i were a synergistic drug combination over a wide concentration range in *BRCA1* mutant cells expressing a low level of EMI1 protein.

It is well established that the “BRCAness” phenotype of the breast cancer cells plays a crucial role toward PARPi sensitivity and conferring the concept of synthetic lethality, and our results are consistent with that concept. Partial restoration of the homologous recombination DNA repair pathway can lead to resistance to PARPi (27), and upregulation of RAD51 in BRCA1-defective cells is associated with resistance to PARPi (20). It is worth noting that our results of olaparib resistance in different BRCA1 mutant cells due to EMI1 downregulation that were associated with high RAD51 levels are in line with previous results done either in BRCA1 mutant cancer cells (20) or BRCA2 mutant ones (25). As expected, our results extended the findings of low EMI1 levels toward modulating the resistance to another PARPi, talazoparib, and strikingly, our results show resistance upon EMI1 downregulation to other cytotoxic drugs including cisplatin and CHK1i. We hypothesized that the RAD51 increase that accompanied the low EMI1 levels may explain the resistance as high RAD51 expression is reported to be associated with decreased cytotoxicity to DNA damage induced by chemical agents and/or ionizing radiation (14,28,29) and RAD51 downregulation can lead to chemo/radio-sensitizing effect (30). However, RAD51 overexpression only partially recapitulated the phenotype of resistance to PARPi due to low EMI1 levels, suggesting that other effects of EMI1 on cell growth also contribute to the PARPi resistance such as the well known requirement for EMI1 function for cells to enter mitosis. Normal EMI1 function inhibits APC/C and also inhibits DNA re-replication, promoting cell proliferation (31). Inactivation of EMI1 leads to premature APC/C activation in G2-phase, interferes with cyclin B accumulation and consequently precludes mitotic entry. As a result, EMI1-depleted cells bypass mitosis and enter cycles of endoreplication (23,24,32). We suggest that *BRCA1* mutated breast cancer cells can develop PARPi resistance by downregulating the *EMI1* gene which could be triggered via p21 (15). Since *EMI1* is an essential gene in vivo (33) and is also required for long-term growth in vitro (34–36), the downregulation of EMI1 must either be transient or retain residual activity sufficient for cell viability. The downregulation of EMI1 arrests cells in G2 phase of cell cycle, tumor cells that remain in G1- or G2-phase longer are less sensitive to PARP-inhibitor-induced cytotoxicity (25). In addition, down regulation of EMI1 stabilizes RAD51 protein levels since EMI1 regulates the ubiquitin-mediated degradation of RAD51 (20). Our results suggest that these combined effects are responsible for the chemotherapy resistance conferred by EMI1 depletion, and overexpression of RAD51 causes only partial resistance to some PARPi.

It had been predicted that combined effects of CHK1i and PARPi would restore sensitivity of cells that had reduced EMI1 expression levels (8,20). Results presented in this study confirm this notion for BRCA1-mutant breast cancer cells. These results are consistent with recently published results for other BRCA1/2 mutant olaparib sensitive and olaparib resistant ovarian cancer cell lines (37). These results suggest a potential treatment strategy toward overcoming PARPi resistance in BRCA1 associated breast cancer cells.

## Acknowledgments

We would like to thank the members working in Parvin lab for invaluable assistance, including, Aleksandra Adamovich, Tapahsama Banerjee, Xue Wu, Kathryn Duncan, Mariame Diabate, Alexander E. Hare and Gregory R. Nagy. This work was supported by grant CA228083 from the National Cancer Institute and Egyptian Ministry of Higher Education and scientific Research (Mission sector).

## Supplemental Tables

***Supplementary table 1. Sequences of siRNAs used in this study***.

***Supplementary table 2. Source of drugs used for proliferation assays***.

***Supplementary table 3. Combination Index (CI) values for actual experimental datapoints***.

A constant-ratio drug combinations 1:1 of olaparib:CHK1 I was analyzed in all cells tested. When analyzing the proliferation of EMI1-depleted cells versus control siRNA transfected cells, the CI values for all concentrations tested were lower than 1, suggestive of a synergistic effect between the two drugs under these conditions.

